# Multinomial-Poisson mixture models reveal unexpected higher density estimates of an Andean threatened bird

**DOI:** 10.1101/298091

**Authors:** D. A. Gómez-Hoyos, O. H. Marín-Gómez, Y. L. Caicedo Ortiz

**Author notes:** Corresponding author: Oscar Humberto Marín-Gómez.

## Abstract

*Multinomial-Poisson mixture models reveal unexpected higher density estimates of an Andean threatened bird*.Distance sampling and repeated counts are important tools to estimate population density of birds with low detectability. Here we use model based approach to assess the population density of a threatened bird, the Multicolored Tanager (*Chlorochrysa nitidissima*). We conducted 144 fixed point counts samplings to record all the individuals of the Multicolored Tanager detected by visual and aural observations from different habitats (forest edge, mature, secondary, and riparian forest), during four months in an Important Bird Area of Central Andes of Colombia. We used spatially replicated counts, distance sampling, and multinomial-Poisson mixture models to estimate the population density of the Multicolored Tanager. Accumulated sampling effort was of 576 repetitions in 144 point counts with 96 h of observation. The Multinomial-Poisson mixture model showed the best fit due low variance of density estimations in comparison to the conventional distance sampling and the spatially replicated counts. Results of this model evidenced a remarkable higher density estimates (1.3 – 2.05 individuals/ha) of the Multicolored Tanager, particularly in mature and secondary forest, as a result of detection correction, instead of sampling effort, by our model based analysis in contrast to index density used in previous studies. We discuss the advantages of model based methods over density indexes in designs monitoring programs of endangered species as the Multicolored Tanager, in order to obtain better and comparable assessment of density estimations along multiple localities.

**Resumen:** *La combinación de modelos multi-nominales y Poisson revelan estimaciones de densidad altas inesperadas en un ave amenazada Andina*. Los muestreos por distancias y los conteos repetidos son herramientas importantes para estimar la densidad de población de aves con baja detección. Aquí utilizamos un enfoque basado en modelos para evaluar la densidad de población de un ave amenazada, la tangara multicolor (*Chlorochrysa nitidissima*). Realizamos 144 muestreos de conteos de puntos fijos para registrar todos los individuos de la tangara multicolor detectados por observaciones visuales y auditivas en diferentes hábitats (borde del bosque, bosque maduro, bosque secundario y bosque ribereño), durante cuatro meses en un Área Importante para la Conservación de las Aves en los Andes centrales de Colombia. Utilizamos conteos replicados espacialmente, muestreos de distancia y la combinación de modelos multi-nominales y Poisson para estimar la densidad de población de la tangara multicolor. El esfuerzo de muestreo acumulado fue de 576 repeticiones en 144 puntos de conteo con 96 h de observación. La combinación de modelos multi-nominales y Poisson mostró el mejor ajuste debido a la baja varianza de las estimaciones de densidad en comparación con el muestreo de distancias y los conteos replicados espacialmente. Los resultados de este modelo evidenciaron una notable estimación de mayor densidad (1.3 – 2.05 individuos / ha) de la tangara multicolor, principalmente en bosques maduros y secundarios, como resultado de la corrección de la detección por nuestro análisis basado en modelos, en lugar del esfuerzo de muestreo, en contraste con los índices de densidad utilizados en estudios previos. Discutimos las ventajas de los métodos basados en modelos sobre los índices de densidad en los diseños de programas de monitoreo de especies en peligro como la tangara multicolor, con el fin de obtener una evaluación mejor y comparable de las estimaciones de densidad a lo largo de múltiples localidades.

## Introduction

Estimating densities is a basic step to evaluate the status of a population. For land birds, the most accurate results of density arise from a combination of different methods as point counts, linear transects, territory mapping of marked individuals, and nest monitoring (Ralph et al., 1995; Bibby et al., 2000). However, intensive sampling and financial resources are required to ensure collecting enough data, hence point counts have become the standard and non-expensive method to access the abundance and density of bird populations around world (Ralph et al., 1995; Bibby et al., 2000). Distance sampling (Buckland et al., 2001) and repeated counts (Royle, 2004) are model based estimations that improve the confidence of the parameters by taking into account the biases among observers, habitat, and site conditions, which are corrected by the detection probability function (Buckland et al., 2001; Norvell et al., 2003; Hutto, 2016). This approach is essential to understand the population trends of Neotropical birds, particularly of those endemic to montane ranges, which exhibit low densities (Jankowski and Rabenold, 2007). However, the population estimates of endangered bird species are still scarce (Kanegae, 2012) or mainly focused on large frugivorous (Kattan et al., 2014, 2015; Denis et al., 2016; González-García et al., 2017; Quiñónez-Guzmán et al., 2017). In other cases, the few available estimations are not corrected by differences in sampling effort or habitat (e.g. Renjifo et al., 2014). This information is crucial to evaluate the conservation status of species with conservation issues.

The Multicolored Tanager, *Chlorochrysa nitidissima*, is an endemic and global endangered species listed as vulnerable due to its small distribution range and population declining (Fierro-Calderón and Johnston-González, 2014; BirdLife International, 2015). This tanager is restricted to montane forests between 900 and 2200 m of the Western and Central Andes of Colombia, and inhabits primary forests, forest edges, and second growth forests (Collar et al., 1992; Hilty and Brown, 2001; Angarita and Renjifo, 2002). The species forages in pairs in the sub-canopy eating fruits of species of *Cordia*, *Miconia*, *Palicourea*, and *Ficus* (Collar et al., 1992), searching larvae in bromeliads (Cuervo et al., 2008), gleaning underside of leaves (Isler and Isler 1987), and joining to mixed-species flocks (Marín-Gómez and Arbeláez-Cortés, 2015). The population density of the Multicolored Tanager is low compared to other tanager species (Collar et al., 1992) as a result of the fragmentation and loss of 79.3% of its habitat (Renjifo et al., 2014). Therefore, population density studies along the distribution range of this tanager are essential to determine its vulnerability and response to habitat disturbance, since it could optimize conservation efforts.

Despite being a colorful bird, the Multicolored Tanager is relatively difficult to detected during point counts sampling, due to its secretive behavior and rapid foraging movements in the canopy (Cárdenas et al., 2007; Fierro-Calderón et al., 2009; Marín-Gómez and Arbeláez-Cortés, 2015). Thus, its lower detectability could be related with sampling efforts, and differences in habitat use, breeding period, and fruit availability (Fierro-Calderón and Johnston-González, 2014). Population estimations based on models are needed to assess these biases, facilitate further comparisons among different studies (Anderson, 2003; Moore and Kendall, 2004), and provide guidelines about its conservation. Here we use multiple estimation methods as spatially replicated counts, distance sampling conventional, and multinomial-Poisson mixture model to compare the population density of the Multicolored Tanager in an important Bird Area of Central Andes of Colombia.

## Material and Methods

### Study area

The *Cañón del Río Barbas-Bremen* Important Bird Area (BirdLife International 2017) is located in the western slope of the Central Andes of Colombia between 1500 and 2100 m asl (Figure 1). The landscape is characterized by low montane forest patches and exotic plantations (*Eucalyptus* sp. and *Pinus patula*) immersed in a pasture grassland matrix (Figure 1). The two largest patches are the *Cañón del río Barbas* (04º42’38’’ N; 75º38’52’’ W) with 790 ha, and the *Reserva Natural Bremen-La Popa* (04º40’27’’ N; 75º37’56’’ W) with 747 ha. These patches have some areas of well-preserved forest located in deep canyons with abrupt topography (BirdLife International, 2017). Details of study area are provided by Gómez-Hoyos et al. (2014).

**Figure 1.**
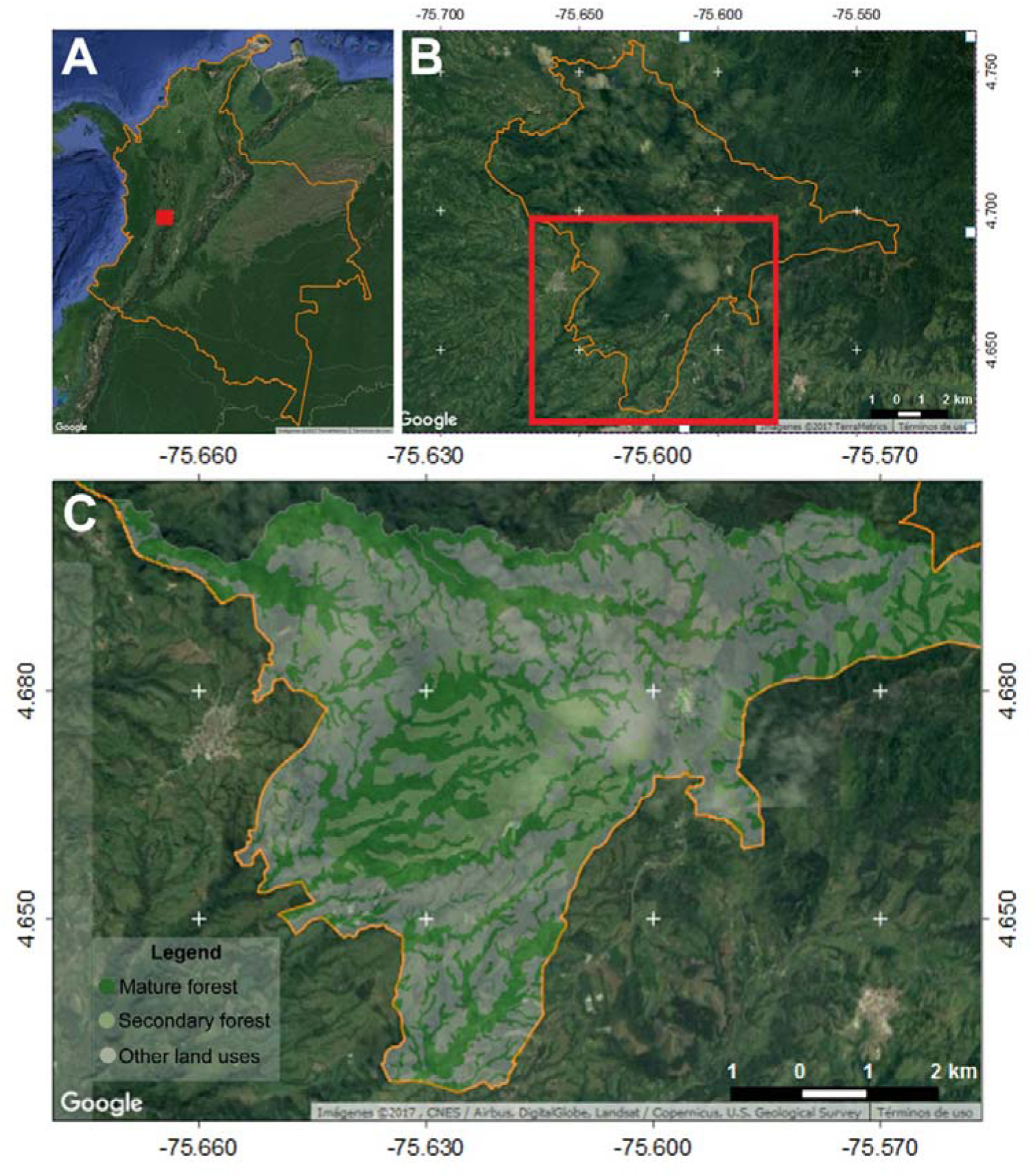
Location of the *Cañón del Río Barbas-Bremen* Important Bird Area (BB IBA) in the Central Andes of Colombia (A), limits of BB IBA (B) and detail of the study area (C). Figura 1. Ubicación del Área Importante para la Conservación de las Aves (AICA- BB) Cañón del Rio Barbas-Bremen en los Andes centrales de Colombia (A), límites del AICA-BB (B) y detalle del área de estudio (C).

### Field sampling

We used fixed point counts of 50 m ratio to estimate the population density of the Multicolored Tanager (Ralph et al., 1995; Bibby et al., 2000). From III to VI 2009, we sampling 144 points placed at intervals of 150 linear meters distributed in four habitat types (Table 1). We sampled each point once per month during four months. Previous to sampling, we marked each counting station using flagging tape at 5 m intervals from the point center towards four cardinal points. Data collection was carried out by two observers (OHMG and DAGH) starting 30 min after local sunrise (06:30) and continued for three hours. Counts were made under similar weather conditions (avoiding rainy and cloudy situations) using 10 x 42 binoculars and a field recorder (Marantz PMD 222 with a Sennheiser ME66) to record any bird sound detected. The observers arriving at each station waited one minute before start counting all the individuals of the Multicolored Tanager detected in a 50 m ratio during 10 minutes. The type of record (aural or seen), time of the first contact, sex, foraging activity, vertical strata, and radial distance were recorded for each encounter. To calculate radial distance we use measure tape from the point center (or the flagging marks intervals) to the place where the bird was detected. In some cases, the exact distance could not be measured, so we assigned the detection to the nearest marked interval.

**Table 1.** Description of the habitat type were the population density of the Multicolored Tanager was studied. Tabla 1. Descripción de los tipos de hábitat donde se estudió la densidad poblacional de la tangara multicolor.

| Habitat type | Description | Canopy height | Dominant tree species | Sampling points |
| --- | --- | --- | --- | --- |
| **Mature forest (MF)** | Well preserved forest remnants located in sharp slopes, with a dense understory | 35 m | *Sphaeropteris quindiuensis*, *Prestoea acuminata*, *Chrysochlamis dependens*, *C. colombiana*, *Gustavia superba*, *Otoba lehmannii*, *Palicourea angustifolia*, *Cecropia telealba*, *Calatola colombiana*, and *Vismia guianensis*. | 24 |
| **Secondary forest (SF)** | Disturbed forest remnants with open understory, pioneer trees and shrub species | 30 m | *Palicourea angustifolia*, *Symplocos quindiuensis*, *Chrysochlamis colombiana*, *Sphaeropteris quindiuensis*, *Ladenbergia oblongifolia*, *Hedyosmum bonplandianum*, *Miconia* sp., *Gustavia superba*, *Alchornea coelophylla*, *Cyathea* sp., *Oreopanax floribundum*, *Ocotea microphylla*, *Axinaea microphylla*, *Miconia lehmannii*, and *Meriania speciosa*. | 67 |
| **Riparian forest (RF)** | Native vegetation patches along the streams with open understory and dominance of herbaceous and shrub species, and some old trees | 20 m | *Meriania speciosa*, *Miconia* sp., *Hedyosmum bonplandianum*, *Geonoma undata*, *Hyeronima scabrida*, *Symplocos quindiuensis*, *Palicourea angustifolia*, *Otoba lehmannii*, *Sphaeropteris quindiuensis*, *Oreopanax floribundum*, *Miconia lehmannii*, *Chrysochlamis dependens*, *Croton magdalenensis*, and *Miconia* sp. | 29 |
| **Forest edge (FE)** | Open understory with herbaceous and shrub species and some old trees | 15 m | *Meriania speciosa*, *Cecropia telealba*, *Miconia lehmannii*, *Croton magdalenensis*, *Symplocos quindiuensis*, *Heliocarpus popayanenses*, *Hedyosmum bonplandianum*, *Montanoa quadrangularis*, *Palicourea angustifolia*, *Chrysochlamis colombiana*, *Miconia* sp., *Hyeronima scabrida*, *Palicourea acetosoides*, *Cupressus lusitanica*, *Oreopanax floribundum*, and *Cordia cilindrostachya*. | 24 |

### Data analysis

We evaluated the performance of repeated counts and distance sampling models to assess the population of the Multicolored Tanager. Mixed models were used for the repeated counts, under the assumption of consecutive visits to the point counts, which were replicated temporally and spatially during the sampling period (Royle, 2004). We chose the upper limit of model integration (k) as 50, which represents an additional unit to the maximum number of individuals detected for point count, multiplied this value by 10 (Wenger, 2008). Since this model estimates the abundance and in order to compare with the other estimates, we calculated the difference between abundance and detection area in each point as pi*r^2^ (3.1416*28^2^), where r is the effective detection radius estimated with the distance sampling method. We used the Poisson distribution due to its best adjustment to count data. Models were generated using the *pcount* function in the *Unmarked* package (Fiske et al. 2015) of R language (R Core Team, 2017).

The distance sampling methods were adjusted to conventional models (Buckland et al., 2001; Thomas et al., 2010), meanwhile the Multinomial-Poisson mixture model was used to evaluate the covariate effects of the habitat type on species density (Royle et al., 2004). Distance sampling conventional models were generated using Distance 6 release 2 (Thomas et al., 2010). The models were based on the Half Normal, Uniform, Hazard rate, and Negative exponential functions in combination with the Cosine, Simple Polynomial, and Hermite polynomial expansion series. The analysis were stratified by habitat type. Finally, the Multinomial-Poisson mixture model (Royle et al., 2004) was adjusted to point counts and the distances generated in discrete intervals using the *distsamp* function in the *Unmarked* package (Fiske et al., 2015). These models included the detection functions described above in combination with a null model for the detection and the abundance, as well as models where these variables are affected by habitat type.

The best-fitting models were selected based on the Akaike Information Criteria with a correction for small sample sizes (AICc) where the values with less AICc indicate the most plausible model (Burnham and Anderson, 2002). The model with the best fit was used to estimate the Multicolored Tanager density and the detection probability. When we found uncertainty about the best fitting model, we reported all estimations of top-ranked models (Delta AICc L 2) according to Arnold (2010).

## Results

The accumulated sampling effort was of 576 repetitions in 144 point counts with 96 h of observation. Thirty-three records of 56 individuals of the Multicolored Tanager, mostly in May and June (30 individuals) were obtained. Most records (73%) were aural, which correspond presumably to pairs. We also detected solitary individuals and conspecific groups conformed by a male, a female, and an immature. Most of visual records corresponded to birds foraging in pairs or conspecific groups in the canopy, or joining mixed flocks (44%).

### Spatially replicated counts

The best fitting model for spatially replicated counts was abundance non-affected by habitat type and detection explained by habitat type (Table 2). The estimate of density was 1.3 individuals/ha (SE=0.62; IC 95% = 0.59 – 2.87) with a detection probability from 0.036 (SE=0.023; IC 95% = 0.01 – 0.12) in secondary forest to 0.11 (SE=0.062; IC 95% = 0.033 – 0.299) in mature forest (Table 2). The second best model included the abundance explained by habitat type with estimates from 0.95 individuals/ha (SE=0.58; IC 95% = 0.29 –3.11) in secondary forest to 2.98 (SE=1.74; IC 95% = 0.95 – 9.37) in mature forest (Figure 2).

**Table 2.**
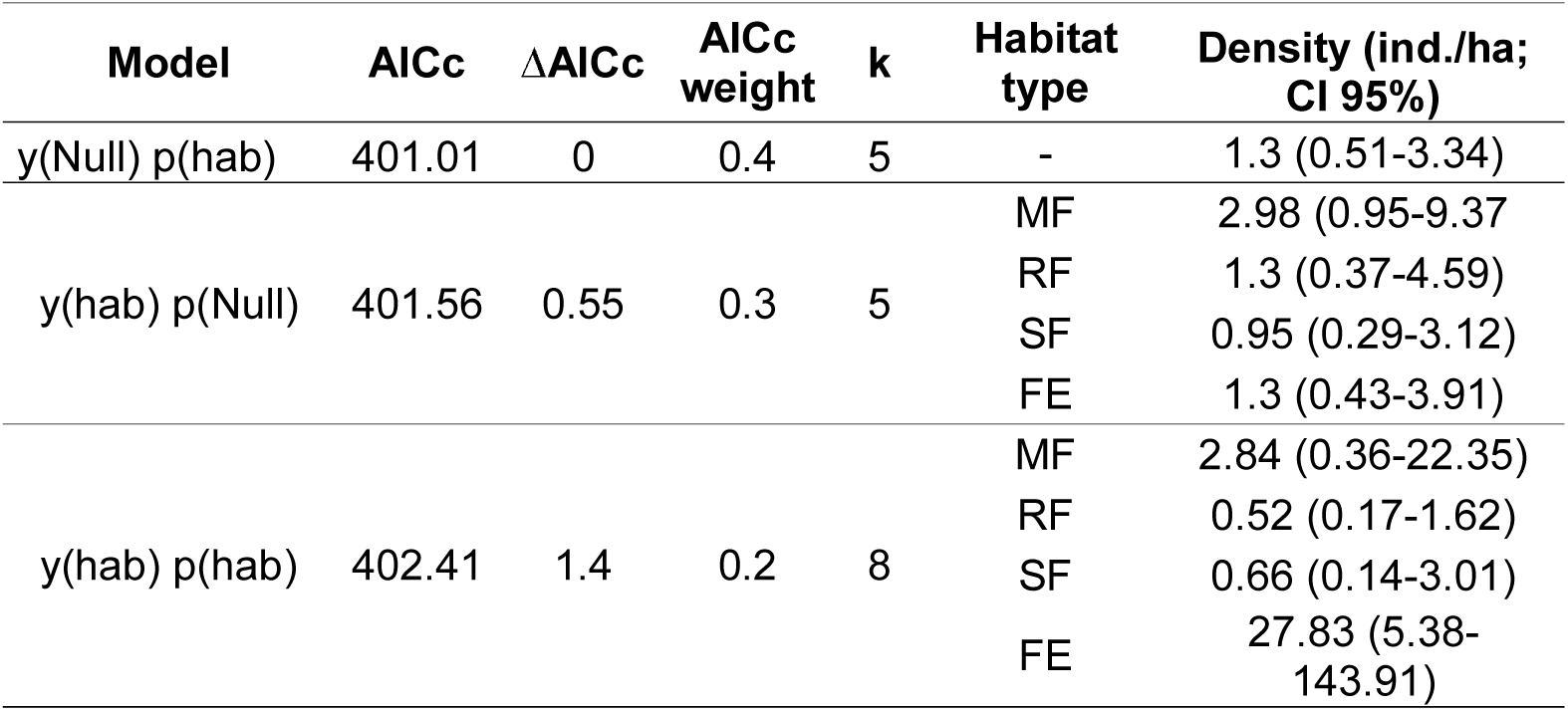
Top-ranked models for N-mixture models and density estimation for the Multicolored Tanager. Tabla 2. Los modelos mejor calificados basados modelos N-mixtos y estimación de la densidad de la tangara multicolor.

| Model | AICc | $\Delta AICc$ | AICc weight | k | Habitat type | Density (ind./ha; CI 95%) |
| --- | --- | --- | --- | --- | --- | --- |
| y(Null) p(hab) | 401.01 | 0 | 0.4 | 5 | - | 1.3 (0.51-3.34) |
| y(hab) p(Null) | 401.56 | 0.55 | 0.3 | 5 | MF | 2.98 (0.95-9.37) |
|  |  |  |  |  | RF | 1.3 (0.37-4.59) |
|  |  |  |  |  | SF | 0.95 (0.29-3.12) |
|  |  |  |  |  | FE | 1.3 (0.43-3.91) |
| y(hab) p(hab) | 402.41 | 1.4 | 0.2 | 8 | MF | 2.84 (0.36-22.35) |
|  |  |  |  |  | RF | 0.52 (0.17-1.62) |
|  |  |  |  |  | SF | 0.66 (0.14-3.01) |
|  |  |  |  |  | FE | 27.83 (5.38-143.91) |

**Figure 2.**
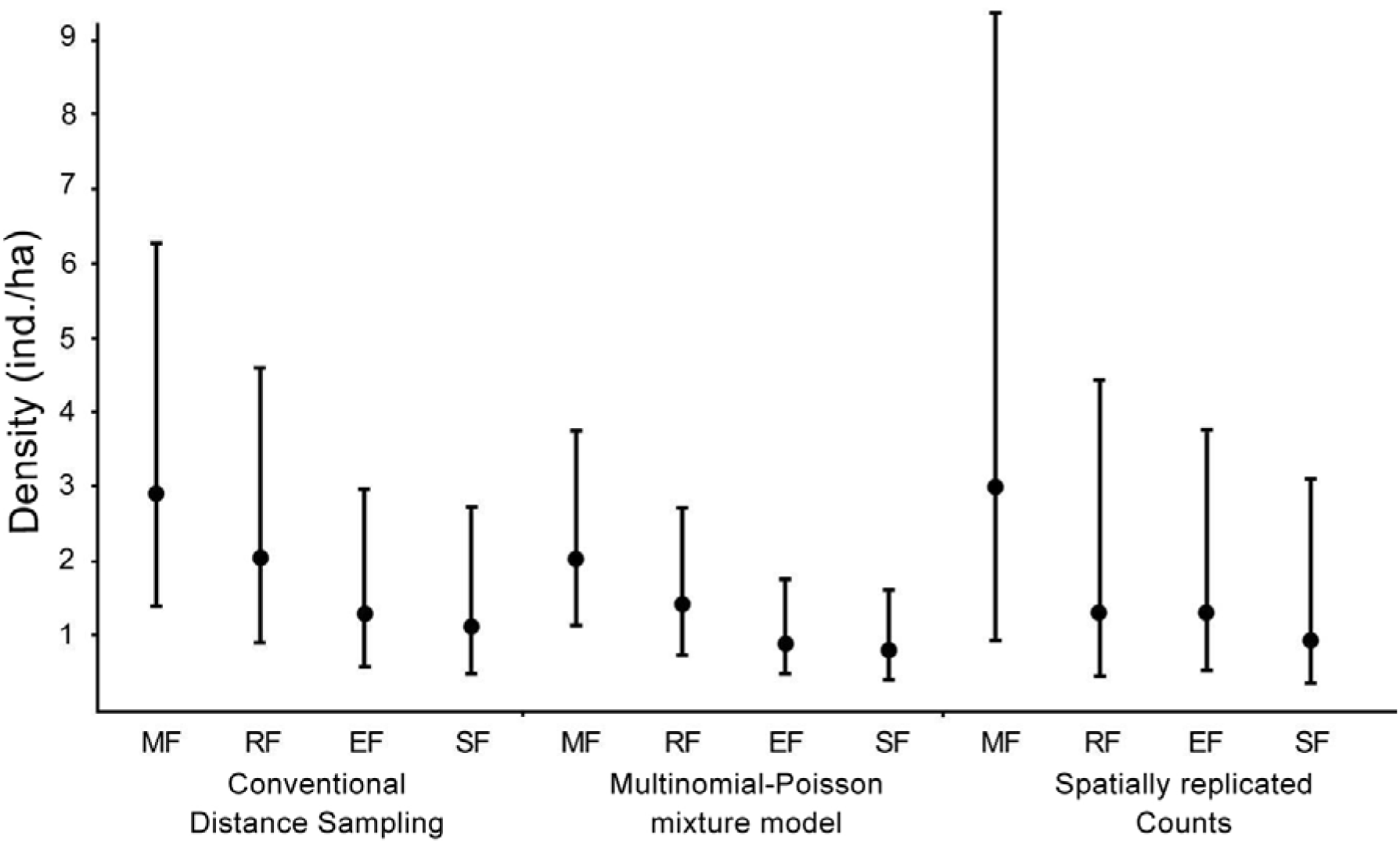
Density estimation for the Multicolored Tanager with different methods by habitat type. MF., Mature forest; RF., Riparian forest; EF., Edge forest; SF., Secondary forest. Error bars: confidence intervals 95%. Figura 2. Estimaciones de la densidad de la tangara multicolor con diferentes tipos de métodos según el tipo de hábitat. MF: Bosque maduro; RF: Bosque ribereño; EF: Borde de bosque; SF: Bosque secundario. Las barras de error indican los intervalos de confianza al 95%.

### Conventional distance sampling

The models that included the Hazard rate function with the three expansion series (Simple Polynomial, Hermite polynomial and Cosine) were the better adjusted to the distribution of radial distances to the point counts with the lower values of AICc (Table 3). The density of the Multicolored Tanager was 1.86 individuals/ha with these models (IC 95% = 1.05 – 3.27; CV = 28.53%; Table 3). The higher density was found in mature forest and riparian forest compared to the forest edges and secondary forest. However, the estimations were not enough accurate to have certainty in the magnitude of the differences on the density among habitat types (Figure 2). The probability of estimated detection was 49.24% (IC 95% = 32.27 – 75.13; CV = 20.94%) with a radial effective detection of 28.01 m (IC 95% = 22.68 – 34.73; CV = 10.47%).

**Table 3.** Top-ranked models for conventional distance sampling and density estimation for the Multicolored Tanager. Tabla 3. Los modelos mejor calificados basados en muestreos de distancia convencionales para la estimación de densidad de la tangara multicolor.

| Function | Series expansion | AICc | $\Delta AICc$ | k | Density (ind./ha; CI 95%) |
| --- | --- | --- | --- | --- | --- |
| Hazard Rate | Hermite polynomial | 229.11 | 0 | 2 | 1.86 (1.05-3.27) |
|  | Simple polynomial | 229.11 | 0 | 2 | 1.69 (0.96-2.95) |
|  | Cosine | 229.11 | 0 | 2 | 1.69 (0.96-2.95) |
| Uniform | Hermite polynomial | 230.32 | 1.2 | 2 | 3.01 (1.46-6.19) |
|  | Simple polynomial | 230.32 | 1.2 | 2 | 3.01 (1.46-6.19) |
|  | Cosine | 231.05 | 1.94 | 1 | 2.77 (1.77-4.35) |

### Multinomial-Poisson mixed models

Based on the AICc values, the models with the best fit included the function Hazard rate with habitat type affecting both the detection probability and species density (Table 4). According to this model, the estimated density for the Multicolored Tanager varied between 2.05 individuals/ha (SE = 1.12; CI 95%: 1.12 – 3.72) in mature forest and 0.79 (SE=0.26; CI 95%:0.45 –1.59) in secondary forest (Figure 2). The highest density was found in mature forest and riparian forest (Figure 2). The detection probability was highest in secondary forest 30.93% (SE=2.96; CI 95%: 26.44 – 36.2), followed by mature forest (24.16; SE=2.99; CI 95%: 19.71 – 29.63), riparian forest (20.36; SE=2.27; CI 95%: 16.95 – 24.47) and forest edge (20.07; SE=1.83; CI 95%: 17.27 – 23.33).

**Table 4.** Multinomial-Poisson mixture models generated for density estimation of the Multicolored Tanager. Tabla 4. Modelos mixtos *Multinomial-Poisson* generados para la estimación de la densidad de la tangara multicolor.

| Models | Function | AICc | ΔAICc | AICc weight | k |
| --- | --- | --- | --- | --- | --- |
| y(hab) p(hab) | Hazard rate | 381.31 | 0 | 0.88 | 9 |
| y(hab) p(null) | Hazard rate | 387.36 | 6.05 | 0.04 | 6 |
| y(null) p(hab) | Hazard rate | 388.12 | 6.81 | 0.03 | 6 |
| y(hab) p(hab) | Half normal | 388.16 | 6.84 | 0.03 | 8 |
| y(hab) p(null) | Half normal | 390.05 | 8.73 | 0.01 | 5 |
| y(null) p(null) | Hazard rate | 390.97 | 9.66 | 0.01 | 3 |

## Discussion

The different methods used here to estimate the population density of the Multicolored Tanager support the low detectability of the species along its distribution (Renjifo et al., 2014). This pattern could be explained by natural history and habitat requirements of this species as it prefers dense cloud forests where forages in sub-canopy and canopy strata, which difficult its observation (Hilty and Brown, 2001; Angarita and Renjifo, 2002). Furthermore, the Multicolored Tanager is vocally active when joining to mixed-species flocks (Marín-Gómez and Arbeláez-Cortés, 2015), where pairs emit constant short contact calls and males sing for few time intervals (Marín-Gómez obs. pers). Hence, aural detections are useful to detect this species.

The Multicolored Tanager is restricted to montane forests from the Western and Central Andes slopes of Colombia (Hilty and Brown, 2001; Angarita and Renjifo, 2002). Although its abundance has been reported higher in the Western than the Central Andes (Renjifo et al., 2014), there are few available density estimates to support this difference. Surprisingly, our results evidenced the opposite, remarkable higher density estimates (1.3 – 2.05 individuals/ha) in a locality of the Central Andes. The available studies using point counts have reported a population density of 0.13 ± 0.16 ind/ha (Fierro-Calderón et al., 2009), and 0.15 ind/ha (Cárdenas et al., 2007). However, those results are probably underestimate, since are based on density indexes which require a constant detection probability, a very difficult task to accomplish (Thompson et al., 1998; Anderson 2001, 2003).

The higher density estimates for this species in our assessment could be related with model based analysis with detection correction in contrast to index density based in the other studies, more than sampling effort. Cárdenas et al. (2007) sampled 80 Km of linear transects for three months, and Fierro et al. (2009) sampled 100 point counts for six months, which is similar to our sampling effort. Including multiple estimation models allows correcting the biases caused by differences in sampling effort and habitat type, which frequently conduct to underestimation of densities.

Distance sampling has been the prevailing method to estimate bird densities, since allows obtaining better estimations in comparison to abundance and density indexes based on count data (Norvell et al., 2003). However, count data are biased by detection errors and zero-inflation affecting inferential power of population status (Dénes et al., 2015). It has been also demonstrated that density estimations using the distance sampling method can reflect the real density of bird populations (Ekblom, 2010). Distance sampling methodology modelling covariate effects assumes that the sampling units are spatially replicated and the distance data are recorded in discrete intervals (Royle et al., 2004). The record of distances in discrete intervals is useful for species as the Multicolored Tanager, as it is difficult to obtain exact measures of perpendicular distances (Ekblom, 2010), breaking one of the assumptions of the conventional distance sampling (Buckland et al., 2001; Thomas et al., 2010). In fact, most obtained records of this study are from vocalizations, which make difficult to take precise measurements of distance, so in these cases, the discrete intervals are recommend (Royle et al., 2004).

Among our assessed models, the Multinomial-Poisson mixture model showed the best fit due to the relatively low variance of density estimations in comparison to the conventional distance sampling and the spatially replicated counts. This model has the advantage of including abundance covariate effects within distance-sampling models (Royle et al., 2004). Besides, when using this model the temporal replication of counts is not needed as is the case of the replicated counts (Kéry et al., 2005, Dénes et al., 2015) with the implication in saving time and budget that it has for designing a monitory program. However, replicated count methods are very competitive when compared to the other rigorous methods for estimating abundance of highly-density species and at large spatial scales (Kéry et al., 2005).

Perfect and constant detectability assumptions are key in a conventional monitoring program using index based estimations (Kéry et al., 2005). Knowing that in our study the Multicolored Tanager detectability was less than 1and heterogeneous among habitats, the perfect and constant detectability assumptions are not ensured. Therefore, to design a monitoring program for this species, it would be necessary to use models to correct by detectability. It is important to point that for low-density species, as is the case of the Multicolored Tanager, it is necessary to increase the spatial representatively of the sampling but not the temporal (in the case of replicated counts) for mitigating the effect of the counts in zero (zero-inflation) and their implications for the inferential capability (Dénes et al., 2015). For all the mentioned before, it is highly recommended to use the distance sampling multinomial-Poisson mixture model to make evaluations or monitoring for the species.

## Acknowledgments

This research was supported by *Fundación Ornitológica del Quindío*. *Corporación Autónoma Regional del Quindío* authorized field work in the *Cañón del Río Barbas- Bremen*. IDEAWILD and Optics for the Tropics provide some equipment. We acknowledge Pedro José Cardona Carmona, Diego Duque Montoya, Diego Martinez, Felipe Carmona, Alba Lucia Lopez, Hernando Castro and Susana Giraldo for the help and companion during the field work. Margarita López García for the comments on an early version of the manuscript. OHMG was supported by the graduate grant 417094 provided by CONACYT. Authors declare that not have conflicts of interest.

